# Modulation of Host Cell Membrane Biophysics Dynamics by Neospora caninum: A Study Using LAURDAN Fluorescence with Hyperspectral Imaging and Phasor Analysis

**DOI:** 10.1101/2024.10.27.620510

**Authors:** Marcela Díaz, Carlos Robello, Andrés Cabrera, Leonel Malacrida

## Abstract

*Neospora caninum* is known to manipulate host cell organelles and recruit lipids for its survival. However, the impact of this lipid redistribution on host cell membranes remains poorly understood. This study used LAURDAN fluorescence, hyperspectral imaging, and phasor plot analysis to investigate how *N. caninum* modifies membrane order in Vero cells. The results revealed a significant decrease in host cell plasma and internal membrane order upon infection, suggesting that cholesterol is redistributed from the host plasma membrane to the parasitophorous vacuoles. To mimic cholesterol depletion, uninfected cells were treated with methyl-β-cyclodextrin (MBCD), which increased membrane fluidity. Conversely, replenishing infected cells with cholesterol-loaded MBCD restored membrane fluidity to levels lower than control cells, indicating cholesterol enrichment. These findings provide novel insights into how *N. caninum* modulates host cell membrane dynamics through lipid manipulation, potentially aiding its intracellular survival.

## Introduction

*Neospora caninum* is the causative agent of bovine neosporosis, a disease responsible for significant economic losses in livestock production worldwide.^1–3^ Despite its global impact, no effective vaccine or treatment currently exists for this parasitic infection, highlighting the need for a deeper understanding of the biology of *N. caninum* and its interactions with the host.^4^ As an obligate intracellular parasite, *N. caninum* relies heavily on the host cell to survive and replicate once the infection is established.^5^ A critical aspect of its life cycle is the invasion process, which leads to the formation of a parasitophorous vacuole (PV), a specialized compartment where the tachyzoite form of the parasite proliferates, shielded from the host’s immune defenses.^6,7^

The PV protects the parasite and serves as a hub for manipulating the host’s cellular machinery. Through this compartment, *N. caninum* initiates signaling pathways that reprogram critical metabolic processes in the host, including targeting organelles like the endoplasmic reticulum, Golgi apparatus, mitochondria, and lipid droplets.^8–13^ Similar to *Toxoplasma gondii, N. caninum* is auxotrophic for cholesterol, relying on the host for this essential lipids. Cholesterol plays a crucial role in membrane structure, making its metabolism a potential target for neosporosis treatment.^8,13–16^ Recent studies have focused on how *N. caninum* alters the host cell’s lipid metabolism, particularly the sequestration of fatty acids and cholesterol into the PV.^13,15^ However, the broader effects of this lipid hijacking on the structure and composition of host cell membranes remain unclear. It is also unknown how these changes create microenvironments within the host cell that differ from those in uninfected cells, particularly regarding membrane order and dynamics.

In this context, LAURDAN (6-dodecanoyl-2-dimethylaminonaphthalene), a solvatochromic fluorescent probe, is commonly used to study membrane dynamics.^17–19^ LAURDAN’s sensitivity to environmental polarity makes it an ideal tool for investigating water dynamics at the membrane interphase, which is significantly impacted by cholesterol content.^20^ In ordered lipid phases, LAURDAN fluoresces at 440 nm, whereas in fluid phases, the emission shifts toward 490 nm.^21–23^

By combining LAURDAN fluorescence with hyperspectral imaging, the whole spectrum can be obtained. Using spectral phasor analysis, LAURDAN fluorescence was demonstrated as a powerful approach for visualizing membrane dynamics.^20,22,23^ The model-free approach and quantitative approximation of the phasor approach enable the address of membrane dynamics in life imaging^24^. By combining these tools with cholesterol depletion or replenishment, we aim to investigate changes in membrane fluidity in *N. caninum*-infected Vero cells. This study will provide new insights into how the parasite alters the host cell’s lipid landscape, which is crucial for understanding its survival mechanisms and potentially identifying new therapeutic targets.

## Materials and Methods

### Cells and parasites cultures

*Neospora caninum* Liverpool strain (ATCC Cat.#: 50845) was obtained from the American Type Culture Collection (ATCC). The strain was cultured as previously described^25^. Briefly, parasites were maintained through serial passages in Vero cells (Vero E6; ATCC # CRL-1586™) using DMEM (Gibco™, Thermo-Fisher Scientific, Waltham, MA, USA) supplemented with 10% fetal bovine serum (Invitrogen, Carlsbad, CA, USA) and 1% penicillin-streptomycin (Invitrogen, Carlsbad, CA, USA). Cultures were incubated at 37°C in a 5% CO_2_ atmosphere. A 1 mL suspension of Vero cells, containing 5 × 10[ cells in DMEM supplemented with 10% FBS, was seeded into T25 culture flask each and incubated at 37°C in a 5% CO_2_ atmosphere for 24 hours. The cells were then infected with purified tachyzoites at a multiplicity of infection of 1:1 in 1 mL of the same medium and incubated again at 37°C in a 5% CO_2_ atmosphere for 96 hours to assess infection. The day before imaging, the infected cell monolayer was detached using 500 μL of 0.25% trypsin. Subsequently, a 1:10 dilution was prepared in DMEM, and 1 mL of this diluted solution was incubated in a 35 mm petri dish with a glass bottom, leaving the plates ready for imaging.

### LAURDAN stain

The membrane probe LAURDAN (6-dodecanoyl-2-dimethylamino naphthalene; Sigma-Aldrich, St. Louis, MO-USA, # 40227) was dissolved in DMSO to make a stock concentration of 3.1 mM. We used 20,000[M−1cm−1 (at 364 nm in methanol) molar extinction coefficient to measure the stock concentration of LAURDAN. Petri dishes were incubated with a 15 µM final concentration in DMEM for 60 minutes at 37°C before imaging.

### Cholesterol depletion

To deplete cholesterol, a 38 mM stock solution of methyl-β-cyclodextrin (MBCD, Sigma-Aldrich, St. Louis, MO-USA, #C4555) was prepared by dissolving it in serum-free DMEM following the protocol of Peres et al^22^. Vero cells were then incubated with MBCD at a final concentration of 10 mM for 60, 90, or 120 minutes at 37°C. After the incubation period, the cells were washed three times with serum-free DMEM and then incubated for 60 minutes at 37°C with LAURDAN dissolved in serum-free DMEM. The dishes were then prepared for imaging.

### Cholesterol enrichment

For these experiments, Vero cells infected with Nc were incubated for 60 minutes with 1 ml of a 2.5 mM MBCD/cholesterol solution (8:1 mol/mol). The cholesterol used (ovine wool cholesterol from Avanti Polar Lipids, INC, #700000) and the MBCD/cholesterol mixture was prepared following a protocol similar to that described by Levitan et al^26^. After incubation, the cells were washed three times with serum-free DMEM. Subsequently, they were incubated for 60 minutes at 37°C with LAURDAN dissolved in serum-free DMEM before imaging.

### Hyperspectral imaging

Spectral imaging was conducted using a Zeiss LSM 880 confocal microscope (Zeiss, Oberkochen, Germany) equipped with a 63x oil immersion objective lens (NA 1.4, Plan-Apochromat, DIC M27, Zeiss). Image acquisition was performed in lambda mode using the Zen Black 2.3 software (Zeiss, Oberkochen, Germany). LAURDAN was excited using a 405 nm diode laser. Emission spectra were captured using a spectral detector equipped with a gallium arsenide phosphide photomultiplier tube (GaAsP-PMT) (650V master gain), collecting data across 30 consecutive channels (figure S1). Spectral images were recorded at a resolution of 512 x 512 pixels, with a spectral resolution of 10 nm, covering a wavelength range of 418 to 718 nm. The pixel dwell time was set to 0.77 μs/pixel, with a pixel size of 146 nm. During live-cell imaging, environmental conditions were carefully maintained at 37°C and 5% CO_2_ to ensure cell viability.

### Phasor plot analysis and interpretation

Hyperspectral data was analyzed using the spectra phasor transformation implemented in SimFCS software, developed by the Laboratory for Fluorescence Dynamics (www.lfd.uci.edu). This approach, which is based on the Fourier transform, processes the spectral data of each pixel in the images as described by Malacrida et al. (2016)^27^. The spectral phasor transform is defined as:

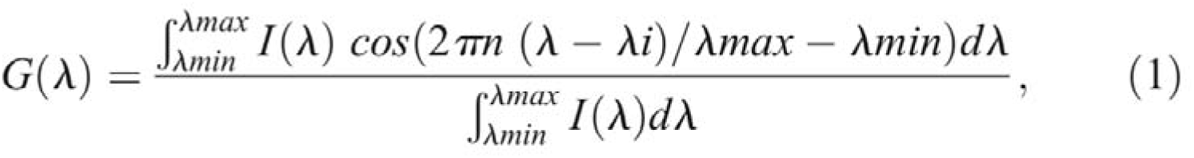

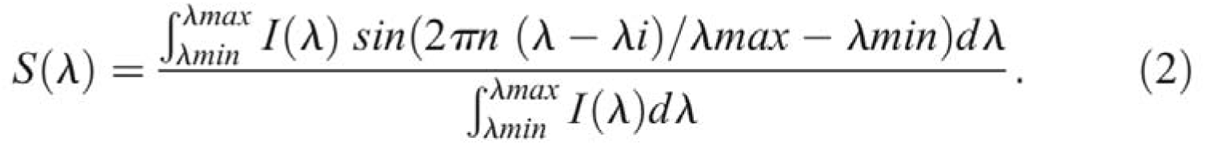

I(λ) is the intensity at each step, n the harmonic number, 1 in our case, λi is the initial wavelength. In the spectral phasor analysis, each image pixel is assigned G and S values, which are then plotted on a polar plot (the spectral phasor plot). This data is visualized as clusters or clouds of points within the four-quadrant phasor plot (2π). Using SimFCS software, the coordinates for each pixel are determined according to equations (1) and (2), mapping each pixel to a specific position in the phasor space. The position on the phasor plot is determined by the spectrum center of mass (phase angle, Ө) and its full width at half maximum (modulation, M). A spectral shift toward longer wavelengths increases the phase angle, while a broader spectrum shifts the point toward the circle’s center (indicating a decrease in M). The spectral phasor approach leverages vector properties, such as linear combination and the reciprocity principle. The linear combination facilitates the analysis of mixtures by representing them as the sum of fractions of individual components. On the other hand, the reciprocity principle enables tracing a region of interest at the spectral phasor plot on the original image (See Figure S1). For all data analysis, denoising of the phasor plots was achieved by applying a 3x3 median filter using SimFCS software.

To analyze the LAURDAN trajectory we defined two individual cursor positions (red and purple) based on the extremes of the pixel distribution for the LAURDAN spectral data in uninfected Vero cells. Lipid membranes exhibit phase transitions, and LAURDAN can change its emission spectrum depending on the phase detected in the lipid bilayer. Its spectrum moves towards green wavelengths in the liquid disorder (L_d_) phase^21^. On the other hand, the LAURDAN spectrum shifts towards blue in the presence of liquid-ordered (L_o_) phases. The terms order and disorder refer to the degree of freedom of the lipid forming the membrane^28,29^. A two-cursor analysis was done using the SimFCS software to quantify the fraction of each component using the addition rules of phasor plots^21^. In this work, fraction histograms were obtained by analyzing the proportions of the components as the linear combination between cursors red (L_d_) and purple (L_o_)^30^. As performed before by Malacrida et al. through the segmentation tool of SimFCS software, we generated masks for individual cells and separated host plasma membranes from internal membranes and, in some cases, PVs^30^. Then, a quantitative analysis was made, measuring the contributions of each component through fractional histograms. In this representation, the number of pixels at each position in the plot is represented along the line or trajectory L_o_ - L_d_, normalized by the total fraction. Then, we could plot the mean curve for each histogram + standard deviation for each point.

We used the Center of mass (CM) calculation for the fraction histograms as a central mean value and the distribution range as dispersion values for statistical

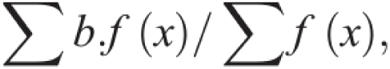

comparison between different groups. The CM was calculated as follows:

With ‘b’ being the percentage of pixels at the fraction ‘f(x)’ of the variable ‘x’, namely purple cursor (L_o_) fraction^31^.

### Statistics

Three independent biological replicates were used to validate this work. For statistical analysis, 8-14 images were acquired per condition. Mean curves for fraction histograms were shown as mean + standard deviation. Center of mass plots were shown as median values for each condition with the dispersion of data. Violin plot was the graphic method chosen; the design was generated through SuperPlotsOfData – a web app for the transparent display and quantitative comparison of continuous data from different conditions^32^. Various conditions were compared using a two-tailed unpaired student t-test with a 95% confidence interval. A One-way ANOVA test was used, followed by Dunnett’s multiple comparisons post-test to compare three groups. All the samples showed normal distributions, as verified by D’Agostino & Pearson, Shapiro-Wilk, and KS normality tests. In all the cases, a p-value [ 0.05 was considered statistically significant. Statistical analysis was performed using GraphPad Software 8, Inc.

## Results

### *N. caninum* infection decreases host cell membrane order

To investigate changes in the lipid composition of host cell membranes, Vero cells were infected with the *Neospora caninum* Liverpool strain (Nc) and stained with LAURDAN (Fig. 1A). Using the phasor plot approach, a significant decrease in membrane order or an increase in overall fluidity was observed. Figure 1A displays representative grayscale images of Vero cells and Vero cells infected with Nc (+Nc). The inset in the figure highlights PVs containing Nc. Figure 1B shows the mean phasor and data distribution for both uninfected Vero cells and those +Nc. The data distribution spans two components, indicated by a red cursor (liquid-disordered, L_d_) and a purple cursor (liquid-ordered, L_o_), defining the fluidity axis. Pseudocolor images in Figure 1C show uninfected cells primarily in green to blue, whereas infected cells shift towards green to red, indicating increased fluidity in the infected cells.

**FIGURE 1:**
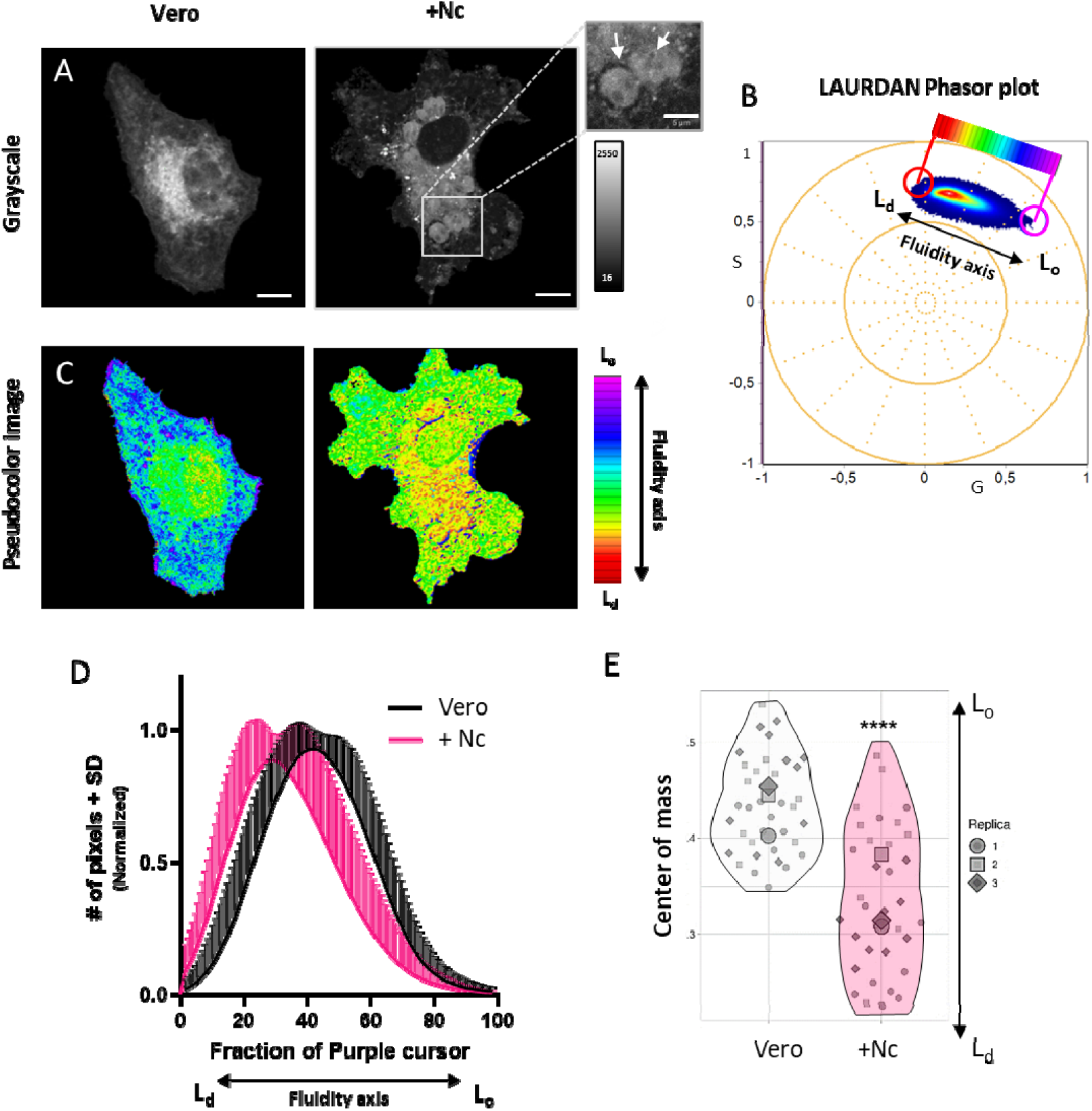
Hyperspectral imaging of LAURDAN fluorescence in uninfected and infected Vero cells with N. caninum. (A) Representative grayscale images of Vero cells and Nc-infected Vero cells (+Nc). The intensity scale (right) is shown in counts, and PV are highlighted (white arrows). Scale bar: 5 µm. (B) Spectral phasor plot of LAURDAN fluorescence in control and Nc-infected cells. Red and purple cursors indicate trajectories for liquid-disordered (Ld) and liquid-ordered (Lo) membranes, respectively. The color scale represents a linear combination of membrane fluidity. (C) Pseudocolor images generated based on the phasor plot in (B). (D) Histograms of normalized pixel counts vs. the fraction of lo membranes (purple cursor) in control and infected cells, with mean curves representing the center of mass distribution. (E) Superplots showing the center of mass distribution from 3 replicates (violin plots) for both cell types. The black arrow indicates membrane fluidity (Ld – Lo). N = 12-14 per replica, scale bar: 10 µm. **** pJ0.0001, unpaired two-tailed t-test, 95% confidence interval.

To quantify this, the purple cursor fraction (L_o_) histograms were generated using SimFCS software. Figure 1D presents the average number of normalized pixels and standard deviation for each linear combination step between both cursors, plotted against the purple cursor fraction. A shift towards the L_d_ phase of the +Nc indicates a decrease in membrane order. Data from three replicates (12– 14 cells each) were analyzed, with centers of mass for each curve calculated (Vero: 0.43 ± 0.05; +Nc: 0.34 ± 0.07). A Superplot in Figure 1E illustrates the means and distribution, showing a 21% reduction in the mean center of mass for +Nc (p-value [0.0001 assessed by unpaired t-test). This result supports a substantial reorganization of cellular membrane order by the increase in membrane fluidity.

### *N. caninum* Infection decreases membrane order in host cell internal and plasma membranes

To address whether *N. caninum* infection affects the lipid order at plasma membranes or intracellular ones, we study its individual contributions by analyzing the phasor of the segmenting region of interest (internal and plasma membranes).

As illustrated in Figures 2A and 2B, an increase in membrane fluidity is observed in both the internal and plasma membranes of +Nc cells compared to uninfected Vero cells. The spectral swift observed by LAURDAN due to Nc Vero infections is visualized in the pseudocolor images, where infected cells display a predominance of green to red hues. On the other hand, bluish-to-greenish colors are identified in Vero cells.

**FIGURE 2:**
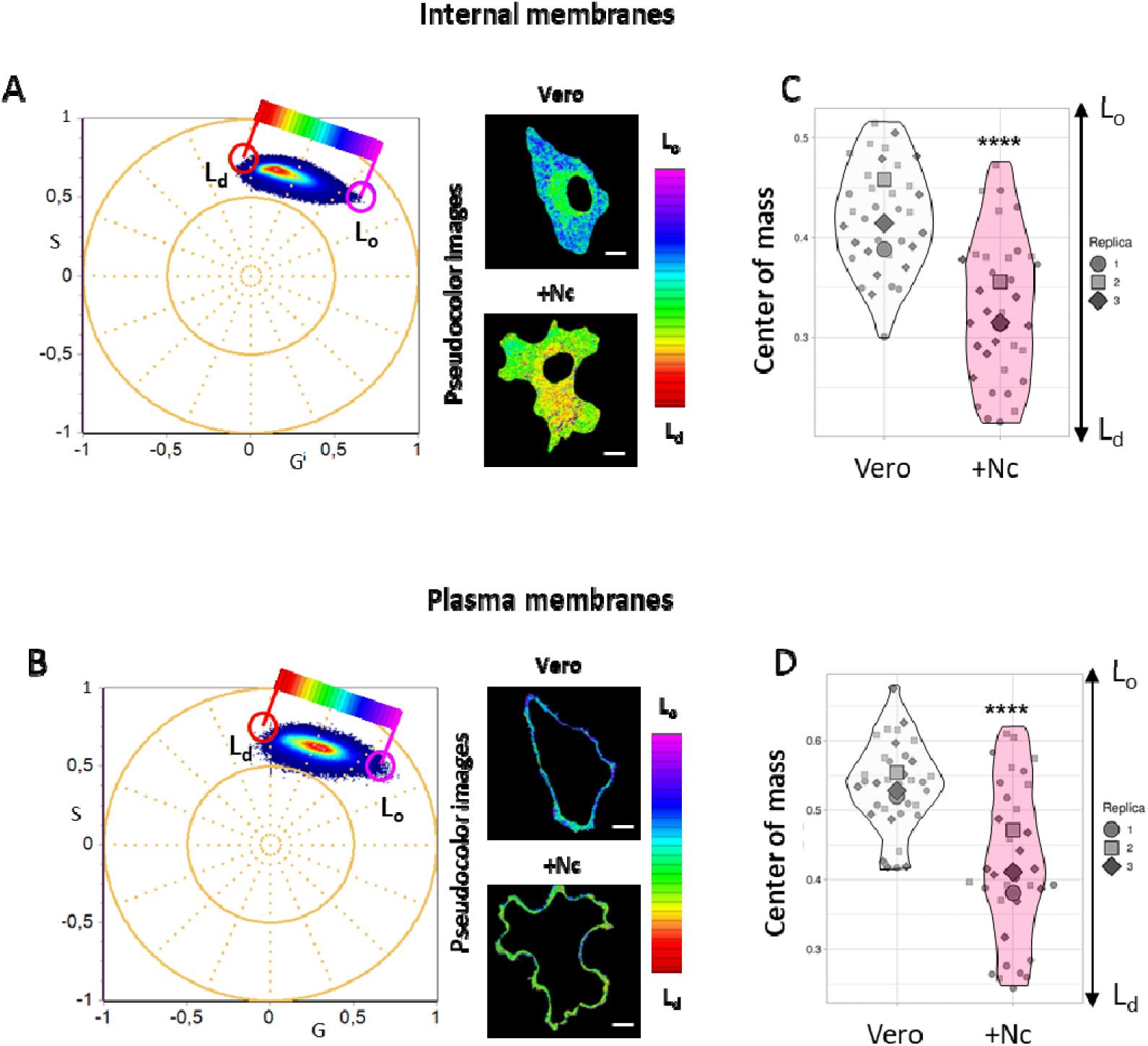
Quantitative LAURDAN-Spectral Phasor Analysis of Plasma and Internal Membrane Fluidity in Vero and +Nc Cells. Spectral phasor plot analysis of LAURDAN fluorescence in internal (A) or plasma membranes (B). The data distribution is shown along the fluidity axis (Ld – Lo) as defined in Figure 1. Pseudocolor images (right) are generated using the red-to-purple color scale. (C, D) Superplots displaying the center of mass of LAURDAN trajectory from 3 replicates (violin plots) for Vero and +Nc cells of internal and plasma membranes respectively. The black arrow indicates membrane fluidity (Ld – Lo). N = 12-14 per replica. **** pJ0.0001, unpaired two-tailed t-test, 95% confidence interval. Scale bar: 10 µm.

Figures 2C and 2D show the superplot analysis of the centers of mass for internal (Fig. 2C) and plasma (Fig. 2D) membranes. The fluidity in the plasma membranes of Vero cells was higher than in internal membranes, and the mean center of mass was 0.53 ± 0.06 vs. 0.42 ± 0.05, respectively. After infection, there was a shifting in the center of mass to 0.33 ± 0.07 for internal membranes. On the other hand, plasma membranes CM shifted to 0.42 ± 0.11 in +Nc cells compared to 0.53 ± 0.06 in control cells. These differences were statistically significant, as confirmed by an unpaired t-test (p < 0.0001).

### MBCD incubation increases the fluidity of host cell membranes depending on concentration and time

*Neospora caninum* can sequester cholesterol^8^. To build on the previous observation of increased membrane fluidity in Nc-infected Vero cells, we studied the LAURDAN response to cholesterol sequestration by MBCD in control cells over time for 60, 90, and 120 minutes. Hyperspectral images analyzed via phasor plot revealed a time-dependent increase in the fluidity of both the internal (Fig. 3A) and plasma membranes (Fig. 3C). This increase was more pronounced at 90 and 120 minutes compared to the 60-minute treatment, as evidenced by the shift from yellow to red hues in the pseudocolor images. Quantitative analysis of the centers of mass for the internal (Fig. 3B) and plasma membranes (Fig. 3D) showed the following values for MBCD-treated Vero cells: at 60 minutes (internal: 0.34 ± 0.05; plasma: 0.52 ± 0.03), at 90 minutes (internal: 0.25 ± 0.03; plasma: 0.41 ± 0.04), and at 120 minutes (internal: 0.23 ± 0.03; plasma: 0.41 ± 0.05). The superplot shows these results together with the average centers of mass for untreated Vero cells (black) and +Nc cells (red), as indicated by the dotted lines. For internal membranes, MBCD treatment significantly increased fluidity at all time points compared to untreated Vero cells (Fig. 3B). In plasma membranes, the increase in fluidity was significant only at 90 and 120 minutes (Fig. 3D). Statistical significance was confirmed using one-way ANOVA followed by Dunnett’s multiple comparisons post-test (p < 0.0001).

**FIGURE 3:**
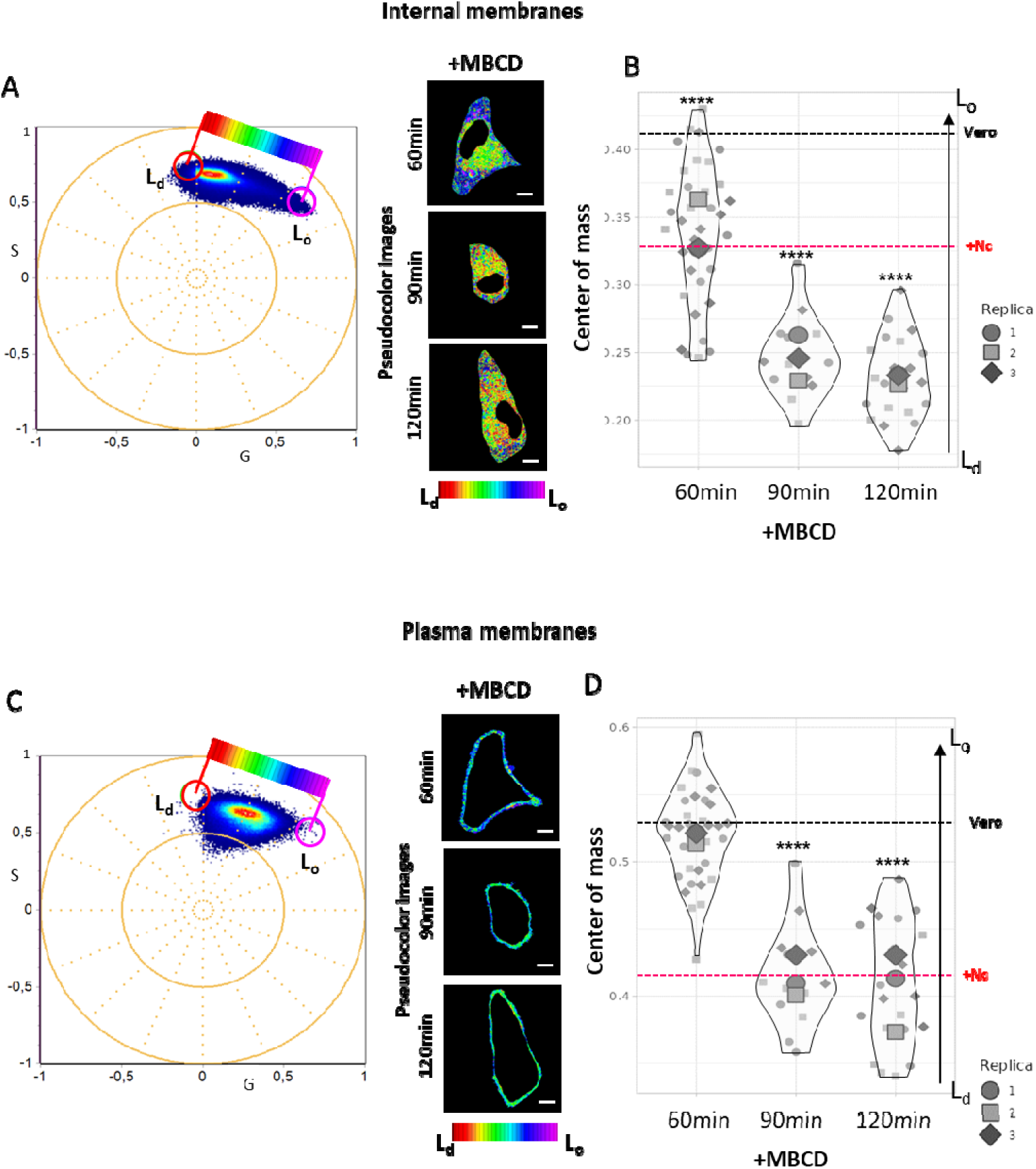
Effect of Cholesterol Depletion by MBCD on Internal and Plasma Membranes Fluidity of Vero Cells. (A, C) Spectral phasor plots of LAURDAN fluorescence in the internal (A) and plasma (C) membranes of Vero cells treated with MBCD for 60, 90, or 120 minutes. Red and purple cursors (Ld – Lo) indicate the boundaries of the data distribution along the fluidity axis. Representative pseudocolor images on the right display fluidity differences, using a color-coded scale, across the different time points after MBCD treatment. (B, D) Center of mass Superplots illustrating the mean and data distribution of internal (B) and plasma (D) membranes from Vero cells treated with MBCD for 60, 90, and 120 minutes. Each replicate (n=3) is represented by a different color. The black dotted line indicates the mean center of mass for uninfected Vero cells (internal: 0.42 ± 0.05, plasma: 0.53 ± 0.06), while the red dotted line represents the mean center of mass for Nc-infected cells (internal: 0.33 ± 0.07, plasma: 0.42 ± 0.11). The black arrow indicates membrane fluidity (ld – lo). N = 5-12 per replicate. ****p < 0.0001, One-way ANOVA followed by Dunnett’s multiple comparison post-test, comparing each MBCD treatment to untreated Vero cells.

### Membrane fluidity restoration in *N. caninum*-infected cells by membrane cholesterol load

To further investigate the role of cholesterol in Nc infection, we introduced cholesterol into infected cells. Exogenous cholesterol was loaded into Vero cells infected with +NcLiv using MBCD-cholesterol (MBCD-chol) for 60 minutes. Figure 4A presents a representative image of MBCD-chol treatment Vero cells infected with Nc. PVs containing Nc are visible in the inset. Phasor plots and pseudocolor images for the internal and plasma membranes are shown in Figures 4B and 4D, respectively. As indicated by the predominance of blue to green colors, MBCD-chol treatment effectively decreased the membrane fluidity of infected cells. Figures 4C and 4E provide quantitative comparisons of the average centers of mass in internal and plasma membranes between +Nc cells treated with MBCD-chol (0.58 ± 0.06, 0.41 ± 0.05, respectively) and untreated +Nc cells. In the superplots, the red dotted line represents the average center of mass for +Nc cells reported in Figure 3. While a significant decrease in membrane fluidity was observed in both membrane types, the reduction was more pronounced in plasma membranes (p < 0.0001) than in internal membranes (p = 0.0004). All statistical analyses were performed using unpaired t-tests.

**FIGURE 4:**
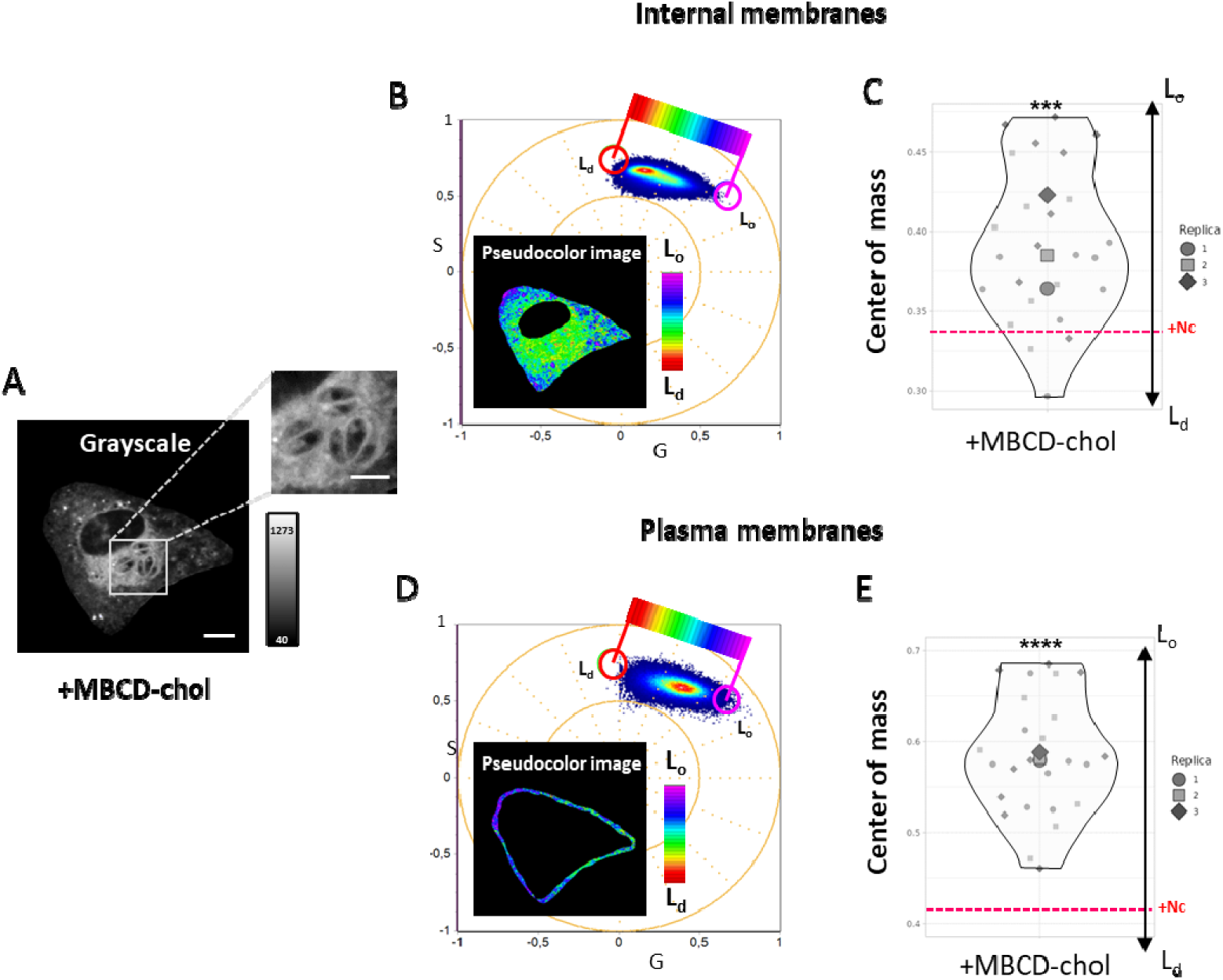
Cholesterol Repletion by Cholesterol-loaded MBCD in Internal and Plasma Membranes of Nc-Infected Vero Cells. (A) Grayscale representative image of a Vero cell infected with Nc and treated with cholesterol-loaded MBCD for 60 minutes (+MBCD-chol). Scale bar = 10 µm (5 µm in the inset). (B, D) Spectral phasor plots of LAURDAN fluorescence in the internal (B) and plasma (D) membranes of +MBCD-chol cells. Red and purple cursors mark the positions corresponding to Ld and Lo membrane phases. The color-coded scale represents membrane fluidity, visualized through pseudocolor images within the phasor plots. (C, E) Superplots displaying the LAURDAN trajectory Center of mass. Mean and data distribution for internal (C) and plasma (E) membranes of +MBCD-chol cells. The black arrow indicates the membrane fluidity range (Ld – Lo). The red dotted line marks the mean center of mass for Nc-infected cells (internal: 0.33 ± 0.07, plasma: 0.42 ± 0.11). Each replicate (n=3) is represented by a different color. N = 8-9 per replicate. ***p = 0.0004, ****p < 0.0001, unpaired two-tailed t-test, 95% confidence interval (+Nc vs. +MBCD-chol).

### Impact of parasite membranes on host cell internal membrane fluidity analysis

To deeper understand the impact of parasite membranes on the observed changes in the fluidity of host cell internal membranes, we examined whether parasite membranes affected the fluidity measurements. Using the image segmentation tool in SimFCS, PVs were isolated in images of Nc-infected cells and cells treated with MBCD-chol. Figures 5A and 5C display the phasor plots and pseudocolor images of the internal membranes of Vero cells without PVs (left) and membranes of PVs (right). Statistical analysis using unpaired t-tests revealed no significant differences between the mean centers of mass for +Nc cells with PVs and those without (0.33 ± 0.08), or between +Nc cells with and +MBCD-chol cells without PVs (0.41 ± 0.05).

**FIGURE 5:**
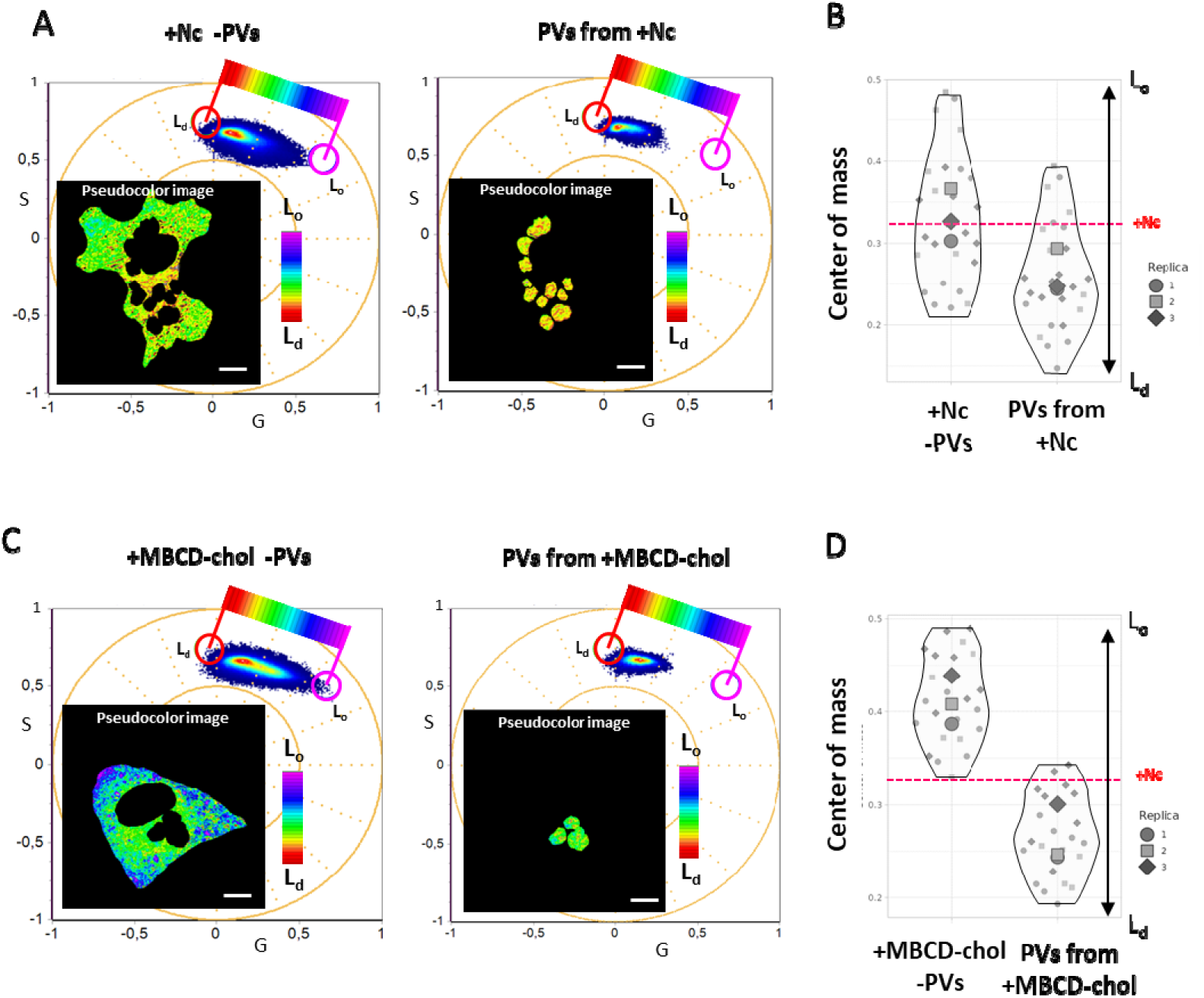
Influence of Parasite Membranes on the overall Host Cell Internal Membranes fluidity. (A) Spectral phasor plots of LAURDAN fluorescence in the internal membranes of Vero cells infected with Nc (+Nc), comparing cells without parasitophorous vacuoles (-PVs) (left panel) and those with segmented PVs (right panel). (C) Spectral phasor plots of LAURDAN fluorescence in the internal membranes of +Nc cells treated with +MBCD-chol, also comparing cells without PVs (left panel) and those with segmented PVs (right panel). Red and purple cursors delineate the fluidity axis from Ld to Lo membrane phases. The pseudocolor images inserted in the plots illustrate the proportion of different fluidity components, as indicated by the color-coded scale. (B, D) Superplots showing the LAURDAN trajectory Center of mass. Mean and data distribution for +Nc (B) and +MBCD-chol (D) cells, comparing -PV and segmented PV conditions. The black arrow represents the membrane fluidity range (Ld – Lo). The red dotted line denotes the mean center of mass for +Nc cells (0.33 ± 0.07). N = 8-12 per replicate (3 replicates per group). No significant differences were observed, unpaired two-tailed t-test, 95% confidence interval (+Nc vs. +Nc -PVs, and +Nc vs. +MBCD-chol -PVs). Scale bar 10 µm.

Superplots for the membranes of +Nc cells (Fig. 5B) and +MBCD-chol cells (Fig. 5D) are presented. In both cases, the PV membranes displayed higher fluidity compared to the internal membranes of host cells, with PV fluidity measurements of 0.26 ± 0.08 for +Nc and 0.27 ± 0.04 for +MBCD-chol.

## Discussion

Understanding the biological mechanisms underlying host-pathogen interactions is crucial for developing strategies to prevent infections^33^. *N. caninum* has been shown to manipulate host cell metabolism and organelles for its own benefit^9–12^. Several studies have focused on how this parasite hijacks and redistributes host cell organelles, particularly mitochondria^10^. Additionally, *N.caninum* is reported to recruit cholesterol and other lipids for its own consumption^8^. However, the impact of this lipid recruitment on host cell membranes and the surrounding cellular environment remains poorly understood.

This study investigates the biophysical changes in membrane dynamics between uninfected and *N. caninum*-infected cells. Using hyperspectral imaging of LAURDAN fluorescence combined with the spectral phasor analysis, we detected an increase in the fluidity of host cell membranes following infection. In normal cells, the literature describes a flow of cholesterol from the endoplasmic reticulum (internal membranes) to the plasma membrane^20,28^. Our findings suggest that infection leads to increased membrane fluidity, potentially due to redirecting the cholesterol path from the plasma membrane to PVs (figure S2).

The global changes induced by infection in the host cell were independently quantified for plasma membranes and internal membranes. This distinction was based on the known differences in cholesterol composition in normal cells, where plasma membranes generally contain a higher proportion of cholesterol^32^. This observation could lead to a differential fluidity between these two cellular environments. However, our findings show that *N. caninum* infection leads to an increase in membrane fluidity in both plasma and internal membranes to a similar extent (figure S2). This result aligns with previous reports indicating that these organisms are auxotrophic, relying on the host cell’s nutrients for survival and replication^8^.

To determine if the absence of cholesterol in uninfected cells could generate an effect on membrane fluidity similar to that observed in Nc-infected cells, we treated uninfected Vero cells with MBCD. MBCD is known to extract cholesterol from plasma membranes without significantly altering phospholipid metabolism after 1 hour of incubation in various cell lines, including Vero cells^22,34^. Phasor plot analysis revealed a significant increase in plasma membrane fluidity 90 minutes after MBCD treatment, while internal membranes showed increased fluidity as early as 60 minutes. Previous studies suggest that even small reductions in plasma membrane cholesterol can cause more extensive cholesterol redistribution in internal membranes^34,35^. This result implies that cells may compensate for cholesterol lost in plasma membranes by redistributing it from internal membranes. Therefore, the observed increase in fluidity is likely correlated with the decrease in membrane cholesterol, potentially affecting other phospholipids after 90 to 120 minutes^34^. After this period, a significant increase in fluidity in both plasma and internal membranes was observed. These findings suggest that cholesterol sequestration by parasites contributes to the increase in membrane fluidity.

Next, we tested whether Nc-infected cells treated with cholesterol-loaded MBCD could decrease fluidity similar to control Vero cells. MBCD can be used to replenish or enrich cells with cholesterol, though the efficiency depends on factors like molar ratios, concentrations, and treatment duration^35^. After 1 hour of treatment, infected cells showed a reduction in membrane fluidity, especially in plasma membranes, with levels lower than those in control cells. This suggests that the treatment is not only replenished but may also have enriched plasma membranes with cholesterol.

For all the phasor plot analyses, parasite membranes were considered together with host internal membranes. As we show in the results of this work, parasite membranes have a high fluidity index, which could introduce bias in the final values. To address this, we performed image segmentation to exclude parasites in their PVs from the final images. Our results confirmed that, in both Nc-infected and MBCD-chol-treated conditions, the parasites did not influence the significant differences in membrane fluidity observed previously.

In conclusion, this study provides important insights into the biophysical changes induced by *N. caninum* infection, specifically in the fluidity of host cell membranes. The observed increase in membrane fluidity across both plasma and internal membranes emphasizes the parasite’s ability to manipulate host cell cholesterol distribution, contributing to its survival and replication. These findings enhance our understanding of host-parasite interactions and offer a foundation for further research into the role of lipid recruitment and membrane dynamics in parasite-host coevolution. By clarifying these mechanisms, this research opens new avenues for developing therapeutic strategies aimed at disrupting these processes and mitigating the impacts of *N. caninum* infections.

