## Supplementary Material for "Modulation of Host Cell Membrane Biophysics Dynamics by Neospora caninum: A Study Using LAURDAN Fluorescence with Hyperspectral Imaging and Phasor Analysis"

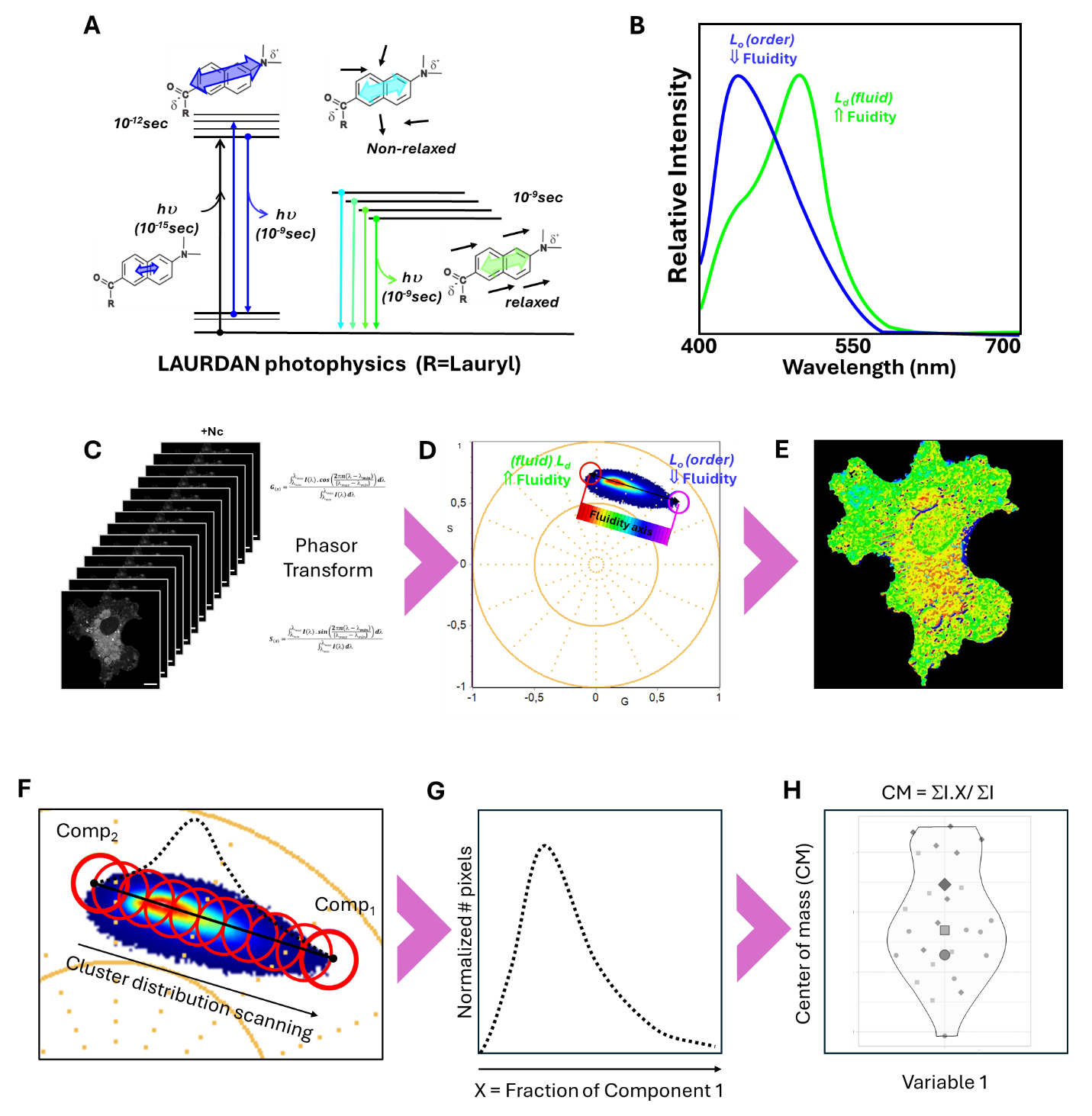

**Figure S1: LAURDAN and spectral phasor rationale analysis.** A) Photophysical properties of LAURDAN. R is a lauryl group in the case of LAURDAN. B) Spectral behavior of LAURDAN fluorescence in liquid-order and fluid (liquid-disorder) membranes. C) Pipeline for hyperspectral imaging analysis using the spectral phasor approach. The hyperspectral stack is transformed into the phasor plot using the G and S formula. A polar plot is obtained, and the cluster is built as a heat map by the accumulation of pixels with similar spectral characteristics. Using the reciprocity principle a color palette is used to identify the different environments experienced by LAURDAN in the cellular membranes. A pseudocolor image is generated, based on the LAURDAN spectral characteristic. F) For the quantitative analysis of the linear combination at the phasor plot, two cursors are used to draw a line that represents the "pure" Lo and Ld membrane and the different fractions in between. Using SimFCS software a histogram of pixel count is constructed. G) The two-component analysis produces a histogram of normalized pixel number and fraction of components. H) to further quantify the difference between the studied group, the center of mass of the histogram distribution is calculated.

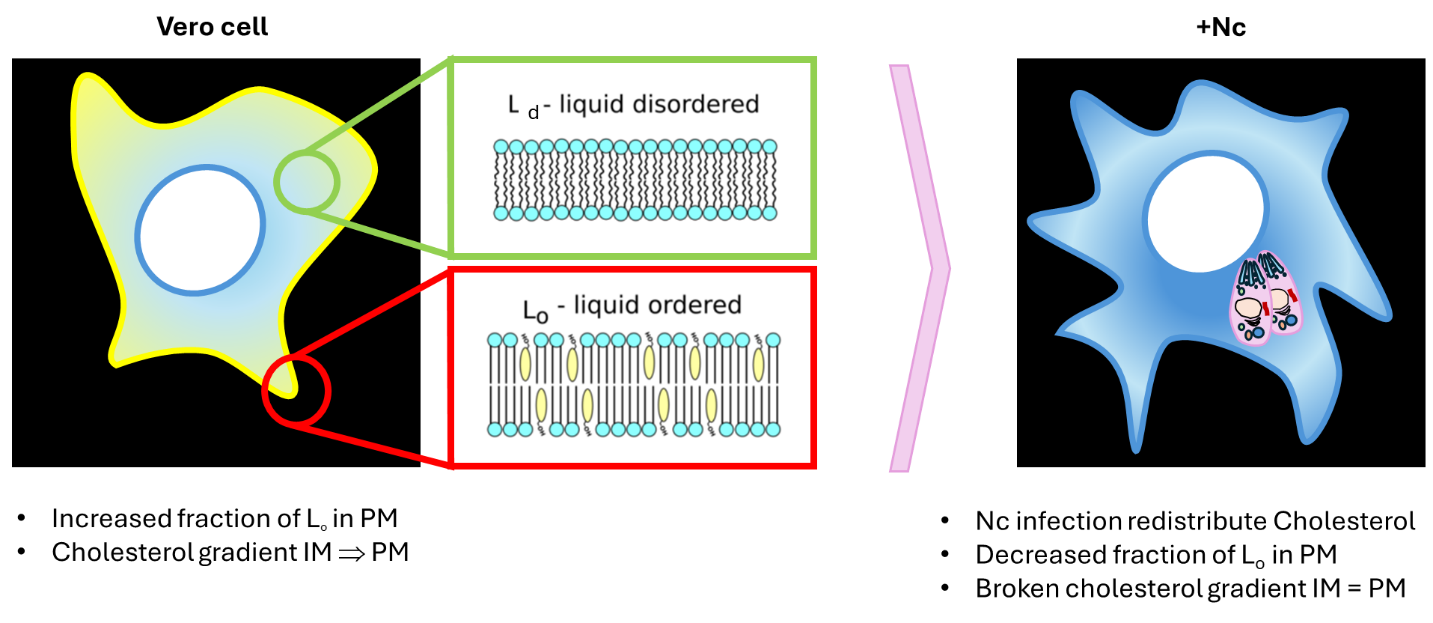

**Figure S2: Model of +Nc infection effects in Vero cell membranes.** Cells have a cholesterol gradient that enables the existence of liquid-order at the plasma membrane and liquid-disorder or fluid at the intracellular interior. Upon Nc infection, Vero cells suffer a redistribution of cholesterol (broken gradient), affecting the membrane order across the entire cells.
